# NEUROMUSCULAR ELECTRICAL STIMULATION LIMITS MUSCLE WEAKNESS, ATROPHY, MODULATES SATELLITE CELL FUNCTION AND REDUCES INFLAMMATION IN CANCER CACHEXIA

**DOI:** 10.64898/2026.04.24.720589

**Authors:** Aliki Zavoriti, Aurélie Fessard, Natacha Boyer, Eugénie Moulin, Camille Koenig, Peggy Del Carmine, Gaëtan Juban, Bénédicte Chazaud, Julien Gondin

## Abstract

**Background:** Cancer cachexia (CC) is characterized by skeletal muscle atrophy and reduced strength, partly linked to dysfunction of muscle stem cells (MuSCs) and alterations in their niche. Although exercise may mitigate muscle loss, its effects in CC remain debated and its feasibility is often limited in advanced patients. Neuromuscular electrical stimulation (NMES) offers a promising alternative, by promoting MuSC proliferation and fusion, increasing muscle size and macrophage content in healthy muscle. This study investigated whether NMES, initiated at tumor onset, could improve MuSC regulation and its niche while limiting muscle atrophy and weakness in a tumor-bearing mouse model.

**Methods:** Ten-week-old male BALB/c mice were subcutaneously injected with C26 tumor cells or PBS. Tumor-bearing mice were divided into NMES-treated (C26 NMES) and non-stimulated controls (C26). NMES consisted of six sessions (two series of three consecutive daily sessions separated by one rest day), starting seven days post-inoculation when tumors became visible. Each session was delivered at a submaximal intensity corresponding to 15% of maximal strength. Muscle mass, myofiber size, strength and cellular composition were assessed.

**Results:** Muscle mass was decreased by 13% in C26 mice as compared to PBS controls, while C26 NMES mice showed a ∼7% improvement over C26 mice. Mean myofiber size decreased similarly in both tumor-bearing groups as compared to PBS controls (−12–14%). However, NMES reduced the proportion of small myofibers (400–600 µm²) as compared to C26 mice. Maximal torque loss was less severe in C26 NMES mice (−28%) than in C26 mice (−34%). As compared with PBS mice, C26 mice exhibited increased MuSC proliferation (+97%) but reduced differentiation (−61%), as indicated by fewer myogenin-positive cells. NMES normalized MuSC proliferation, restored myogenin-positive cell number, and enhanced MuSC fusion, reflected by an increased number of PCM1-positive myonuclei (+8-11%). NMES also modulated inflammation, reducing neutrophils (−42%) and increasing macrophages (+35%), through the proliferation of CD169-positive resident macrophages (+106%). *In vitro*, macrophages exposed to C26 muscle extracts showed elevated pro-inflammatory markers (COX2 and TNF-⍺; +21% and +16%) as compared to PBS controls. This effect was abolished with extracts from C26 NMES muscles. Additionally, C26 extracts reduced the expression of anti-inflammatory markers by macrophages (CD206 and IL-10; −23%), whereas NMES restored their levels to those of controls.

**Conclusion:** NMES-induced mild contractile activity is an effective stimulus for preserving muscle strength and mass, improving MuSC regulation, and modulating muscle inflammation in a mouse model of CC.

## Introduction

Cancer cachexia (CC) is a multifactorial syndrome characterized by an involuntary weight loss that cannot be entirely reversed by conventional nutritional support^1^. It affects over 80% of cancer patients^2^, particularly those with liver or pancreatic cancer, with a one-year mortality rate ranging from 20 to 60%^3^. CC is marked by a severe reduction in skeletal muscle mass (*i.e.,* muscle atrophy), with or without concomitant loss of fat mass^1^, and by functional impairments such as a progressive decline in muscle strength (*i.e.,* muscle weakness) that may occur independently of atrophy^4^. This syndrome profoundly compromises patients’ quality of life, response to chemotherapy and overall survival^5^. Although recent therapeutic approaches, such as ghrelin receptor agonists^6^ or GDF-15 inhibitors^7^, have shown promise, no globally approved treatments currently exist that efficiently prevent or reverse CC.

CC–induced skeletal muscle dysfunctions are primarily attributed to intrinsic myofiber mechanisms, including impaired proteostasis and mitochondrial defects^8^. However, emerging evidence indicates that muscle atrophy during CC also arises from dysregulations of muscle stem cells (MuSCs). MuSCs reside beneath the basal lamina of the myofiber and are critical for muscle homeostasis and regeneration^9^. Upon muscle injury, quiescent MuSCs become activated, proliferate, differentiate into myocytes and eventually fuse with each other or with the existing myofibers to repair damaged myofibers and give rise to new functional contractile units^10^. Interestingly, tumor factors and the systemic inflammation disrupt MuSC regulation as evidence by an increased MuSC number, an impaired differentiation and fusion in both tumor-bearing animals^11–16^ and cancer patients^11,17^.

MuSC regulation also depends on dynamic and tightly coordinated interactions with other cell types within their niche, including immune cells, fibro-adipogenic progenitors (FAPs) and endothelial cells (ECs)^9^. A growing body of evidence reports accumulation of immune cells (e.g., neutrophils, macrophages)^15,18–22^ and FAPs^14^ in skeletal muscle of tumor-bearing mouse models and cancer patients. Collectively, these findings suggest that impaired myogenesis, driven by disrupted cellular interactions within the MuSC niche, play a critical role in the development of CC^21^.

MuSC fate and their niche are also tightly regulated by muscle contractile activity^23^. In response to increased mechanical load or voluntary exercise, MuSCs contribute to myofiber hypertrophy by fusing with myofibers and providing additional myonuclei, a process known as myonuclear accretion^24,25^. Macrophages and FAPs are also essential for muscle growth, actively supporting MuSC fusion and facilitating extracellular matrix remodeling in hypertrophied muscles^26^. Despite this, the effects of exercise on skeletal muscle mass in both tumor-bearing mice and cancer patients are still debated^27–31^. These discrepancies may reflect variations in exercise modality (e.g., wheel running *vs.* treadmill), frequency, duration, intensity, and intervention timing. Notably, beneficial effects are most frequently observed when exercise is initiated prior to tumor cell inoculation or tumor growth, suggesting a predominantly preventive rather than therapeutic role against CC^32^. Moreover, voluntary exercise is often impractical for advanced cancer patients due to poor exercise tolerance, underscoring the need for alternative strategies to enhance muscle activity in CC. In this context, neuromuscular electrical stimulation (NMES), which consists in increasing contractile activity through electrical pulses applied to the skin, emerges as a promising approach. We recently demonstrated that individualized and standardized NMES training protocols delivered at a submaximal intensity promote MuSC fusion and increase both macrophage content and myofiber size in healthy muscles^33^.

Building on these findings, this study investigated whether initiating NMES training at the onset of visible tumor development can improve MuSC regulation and its niche, while minimizing muscle weakness and atrophy in a tumor-bearing mouse model.

## Materials and Methods

### Animals

Ten-week-old BALB/c male mice were purchased from Janvier-Labs (Le Genest-Saint-Isle, France) and were housed individually in a humidity- and temperature-controlled facility (12:12-h light cycle, 25°C). All animals had *ad libitum* access to food and water, except PBS mice, which were pair-fed to C26. Animals were acclimatized to the animal facility during one week before starting the experiments.

### Experimental design

Mice were inoculated subcutaneously in the right flank under general anesthesia with either 5 × 10⁵ C26 tumor cells suspended in 100 µL of PBS (tumor-bearing group) or 100 µL of PBS alone (PBS control group). Tumor-bearing mice were then randomized into two groups: a neuromuscular electrical stimulation (NMES) intervention group (C26 NMES) and a non-stimulated tumor-bearing control group (C26) (Fig. 1A). The NMES protocol consisted of two bouts of three consecutive daily sessions, separated by a single rest day, for a total of six sessions^33^. NMES was initiated seven days post-inoculation, coinciding with the onset of visible tumor growth. The PBS and C26 groups served as non-stimulated controls. Food intake, body weight and maximal torque (T_max_) were recorded two days before injection and from days 7 to 14 post-injection for all groups. On day 14 post-injection, all mice were euthanized by cervical dislocation under deep isoflurane anesthesia. Tumors were harvested and weighed immediately. The *gastrocnemius* muscles were also collected, weighed, and processed for histological or flow cytometry analysis.

**Figure 1:**
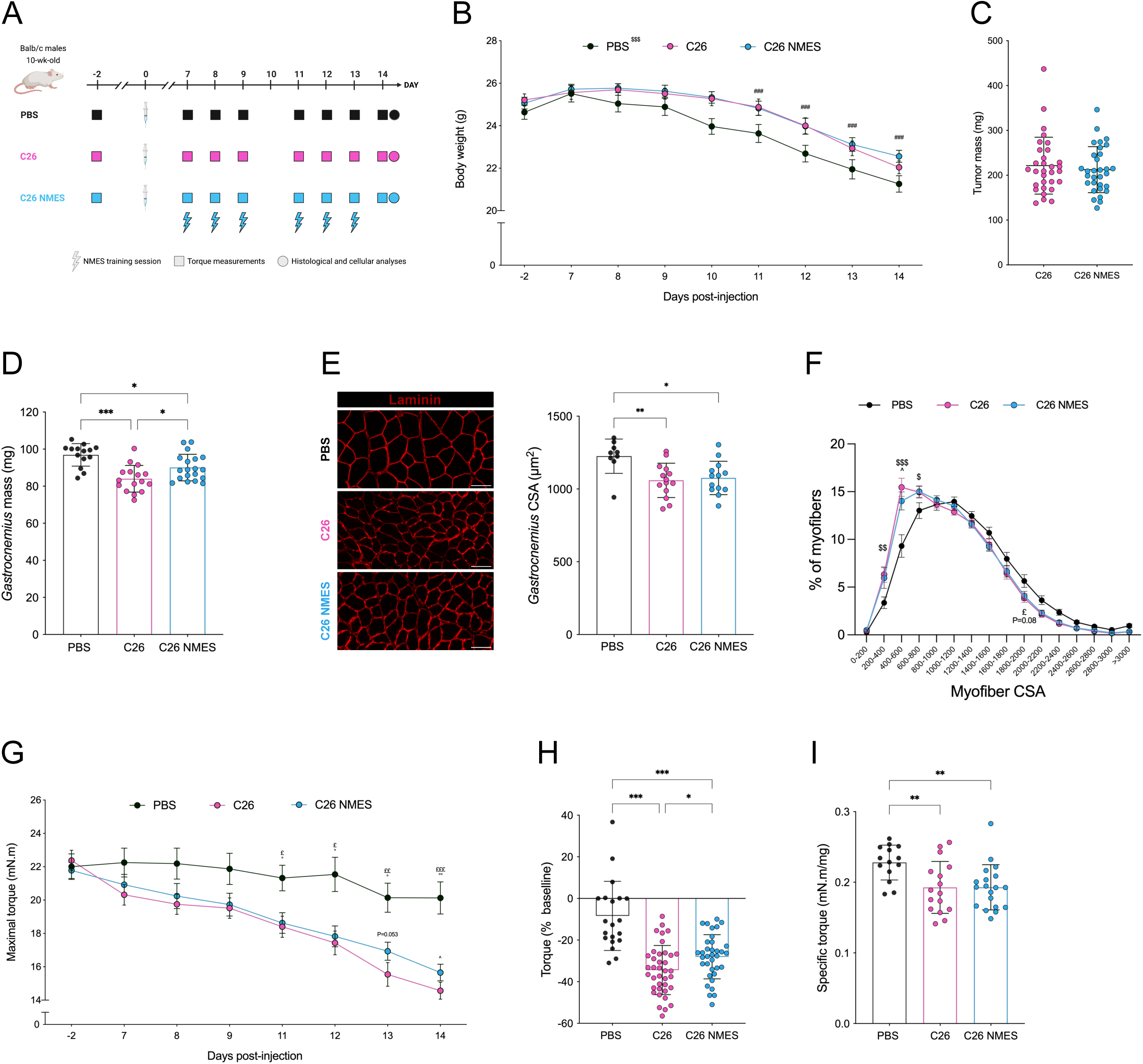
NMES training minimizes loss of muscle mass and torque in the C26 tumor-bearing mouse model without altering tumor mass. A) Experimental design. B) Body weight in PBS (n= 19), C26 (n=38) and C26 NMES (n=35) mice. C) Tumor mass in C26 (n= 31) and C26 NMES mice (n=31). D) *Gastrocnemius* muscle mass in PBS (n=14), C26 (n=16) and C26 NMES (n=19) mice. E) Representative immunostainings and *gastrocnemius* myofiber cross-sectional area in PBS (n=8), C26 (n=8) and C26 NMES (n=9) mice. F) *Gastrocnemius* myofiber cross-sectional area distribution in control PBS (n=9), C26 (n=14) and C26 NMES (n=13). G) Maximal torque production recorded in PBS (n=20), C26 (n=38) and C26 NMES (n=35) mice. H) Changes in maximal torque production from day-2 to day 14 in PBS (n=20), C26 (n=38) and C26 NMES (n=35) mice. I) Specific torque (*i.e.*, maximal torque normalized to *gastrocnemius* muscle mass) in PBS (n=14), C26 (n=16) and C26 NMES (n=19) mice. ^###^Different from day 7 for all groups: P<0.001. ^*,**, ***^P< 0.05, **P< 0.01, P< 0.001. ^$,$$,$$$^PBS different from both C26 and C26 NMES: P< 0.05, P< 0.01 and P<0.001. ^^^C26 NMES different from C26: P< 0.05. ^£,££^PBS different from C26: P< 0.05 and P< 0.01. PBS different from C26 NMES: P=0.08. ^°,°°^PBS different from C26 NMES: P< 0.05 and P< 0.01. Values are reported as mean ± SD (except for panels B, F and G: mean ± SEM).

### C26 cell line

Colon C26 cells were cultured in standard medium (DMEM high glucose, 31966021, Gibco) supplemented with 10% fetal bovine serum and 1% penicillin-streptomycin (15140122, Gibco), and maintained in a humidified incubator at 37°C with 5% CO₂. Cells were passaged at sub-confluence and injected into mice assigned to the tumor-bearing group at passage three.

### Torque measurements and NMES protocol

Mice were placed in a non-invasive ergometer (NIMPHEA_Research, AII Biomedical SAS, Grenoble, France) to electrically stimulate plantar flexor muscles and measure torque production, as previously described. Mice were anesthetized with 4% isoflurane, and the right hindlimb was shaved and prepared with electrode cream. While positioned supine, mice were kept anesthetized with 1.5–2.5% isoflurane administered through a facemask. Electrical stimulation was delivered through surface electrodes near the popliteal fossa and Achilles tendon. The right foot was immobilized on a pedal connected to a torque transducer.

Single twitch stimulation was applied using a constant-current stimulator (Digitimer DS7AH, Hertfordshire, UK; maximal voltage: 400 V; 0.2 ms duration). The current was increased until peak twitch torque plateaued to determine maximal current intensity, which was used to measure T_max_ *via* a 250-ms, 100 Hz tetanic train.

At the start of each NMES training session, stimulation intensity was carefully adjusted to achieve 15% of T_max_. Each NMES session consisted of 80 stimulation trains (2-s duration, 8-s recovery) delivered at a frequency of 50 Hz. Six sessions were performed, yielding a total muscle contractile activity of 16 min (6 × 80 × 2 s). To minimize muscle fatigue and maintain torque at 15% of T_max_ throughout the protocol, the stimulation intensity was increased every 10 contractions^33^.

### Tissue preparation and *in vivo* immunofluorescence analysis

*Gastrocnemius* muscles of the right limb were dissected, frozen in liquid nitrogen-cooled isopentane and kept at −80°C until use. Cryosections (10 µm) were prepared for immunohistochemical analysis with a NX 50 cryostat. A complete description of immunofluorescence protocol is provided in supplemental materials.

#### Muscle protein extracts

A complete description of the experimental protocol is provided in supplemental materials.

#### Myogenesis assessment of MuSC directly exposed to muscle protein extracts

A complete description of the experimental protocol is provided in supplemental materials.

#### BMDM cultures

A complete description of the experimental protocol is provided in supplemental materials.

### Flow cytometry analysis

A complete description of macrophage isolation is provided in supplemental materials. CD45^pos^ cells pre-incubated with FcR Blocking Reagent were stained with AF647-conjugated anti-CD64 (139321, BioLegend), PE-conjugated anti-Ly6C (128007, Biolegend) and BV421-conjugated anti-CCR2 (150605, Biolegend) antibodies. Cells were analyzed on a FACS Canto II apparatus (BD Biosciences). Proportion of CD64^pos^, CD64^neg^Ly6C^pos^, CD64^pos^Ly6C^pos^, CD64^pos^Ly6C^int^ (*i.e.*, expressing intermediate levels of Ly6C), CD64^pos^Ly6C^neg^ were calculated among the total CD45^pos^ cells after analysis by flow cytometry with BD FACSDiva^™^ Software.

#### Image capture and analysis

A complete description of image capture and analysis is provided in supplemental materials.

### Statistical analysis

All statistical analyses were conducted using GraphPad Prism Software (version 10). Normality of data was assessed using the Kolmogorov-Smirnov test. Differences between C26 and C26 NMES groups were evaluated using either an unpaired Student’s *t*-test or Mann-Whitney test. For comparisons among PBS, C26 and C26 NMES groups, either a one-way ANOVA or the Kruskal-Wallis test was applied, followed by Holm-Šídák or Dunn’s post-hoc tests, respectively. Body weight and torque production were analyzed using a two-way repeated measures ANOVA, with Holm-Šídák post-hoc corrections. Data are presented as mean ± SD, except for certain figures where mean ± SEM is reported. Statistical significance was set at p < 0.05.

## RESULTS

### NMES limits muscle loss and weakness in C26 mice independently of tumor growth

To assess the effect of increased contractile activity on CC, we used a standardized NMES protocol validated in healthy mice^33^. Ten-week-old male BALB/c mice were inoculated with C26 tumor cells or PBS. Tumor-bearing mice were randomized to NMES or no treatment. NMES consisted of six sessions in two blocks of three consecutive days separated by one rest day, starting 7 days post-inoculation when tumors were palpable (Fig. 1A). PBS and untreated C26 mice served as controls, and PBS mice were pair-fed to match tumor-bearing mice from day 7.

As previously described^33^, NMES parameters were individualized by adjusting the stimulation intensity to elicit 15% of T_max_ (13.9 ± 2.7% of T_max_; individual mean values ranging from 11.3% to 16.8% of T_max_; Supplemental Fig. 1A). The application of NMES at this submaximal torque level was sufficient to induce a mild contractile activity while avoiding muscle damage, as indicated by the negligible proportion of IgG-positive myofibers (*i.e.*, <1%) across all experimental groups (Supplemental Fig. 1B).

We first evaluated the effect of NMES on body weight. All three groups exhibited a progressive reduction in body weight, starting from day 11 and reaching a loss of 15–18% from their peak values. Because of the pair-feeding strategy, PBS-treated mice exhibited slightly greater body weight loss than C26 tumor-bearing mice, whereas NMES had no significant impact on this parameter (Fig. 1B). We then investigated whether NMES influenced tumor growth. Tumors were harvested and weighed on day 14 post-inoculation. Tumor mass was comparable between C26 and C26 NMES groups, indicating that NMES did not affect tumor burden (Fig. 1C).

Given that CC is characterized by muscle atrophy^34^, we evaluated whether NMES could improve muscle mass and/or myofiber size. *Gastrocnemius* muscles were harvested and weighed on day 14 post-inoculation. Muscle mass in C26 tumor-bearing mice was significantly reduced by 13% compared to PBS controls. Interestingly, muscle mass in C26 NMES mice was ∼7% higher than in C26 mice, suggesting that NMES partially minimized *gastrocnemius* muscle mass loss (Fig. 1D). Myofiber cross-sectional area (CSA) was then determined using immunostaining for laminin (Fig 1E). The mean myofiber CSA was similarly reduced by 12-14% in both C26 and C26 NMES mice compared to PBS controls (Fig 1E). Additionally, the distribution of myofiber size shifted to the left in both C26 and C26 NMES groups relative to PBS (Fig. 1F). Notably, the C26 NMES group exhibited a significantly lower proportion of small myofibers (400-600 µm^2^) compared to the C26 group (Fig. 1F). This indicates that the NMES-induced increase in muscle mass was associated with a mild increase in myofiber size.

Considering that muscle weakness is a hallmark of CC^34^, maximal torque (T_max_) was measured prior to PBS or C26 cell inoculation and subsequently from day 7 to day 14 (Fig. 1A). In C26-bearing mice, T_max_ began to decline from day 7 post-injection (Fig. 1G), preceding body weight loss (Fig. 1B). At 14 days post-injection, the reduction in T_max_ was significantly attenuated in C26 NMES (−28 ± 11%) relative to C26 mice (−34 ± 12%) (Fig. 1G-H). This better preservation of strength was closely linked to increased muscle mass, as specific torque did not differ between C26 and C26 NMES mice (Fig. 1I). Overall, these results demonstrate that NMES effectively limited muscle wasting and weakness independently of tumor growth.

### NMES reduces MuSC proliferation while enhancing MuSC fusion *in vivo*

CC is characterized by dysregulation of MuSCs^11–16^. We recently demonstrated that increased myofiber contractile activity by NMES enhances MuSC proliferation and fusion in healthy mice^33^. Building on these findings, we sought to determine whether the beneficial effects of NMES on muscle strength and mass are associated with improved MuSC regulation. Cryosections from all groups were first immunostained for laminin and Pax7, a marker of MuSCs (Fig. 2A). Consistent with previous studies^11,16^, the number of Pax7-positive cells was 41% higher in C26 tumor-bearing mice as compared to PBS controls (Fig. 2A-B). Importantly, NMES restored Pax7-positive myofiber counts in C26 mice to levels comparable with PBS (Fig. 2A-B). Using immunostaining for Ki67, we further assessed MuSC proliferative capacity. The number of Pax7^pos^Ki67^pos^ cells was 97% higher in C26 tumor-bearing mice relative to PBS controls, highlighting a dysregulation of MuSC proliferation during CC (Fig. 2A&C). Strikingly, NMES completely abolished this aberrant proliferative response, restoring MuSC behavior to levels comparable to controls (Fig. 2A&C).

**Figure 2:**
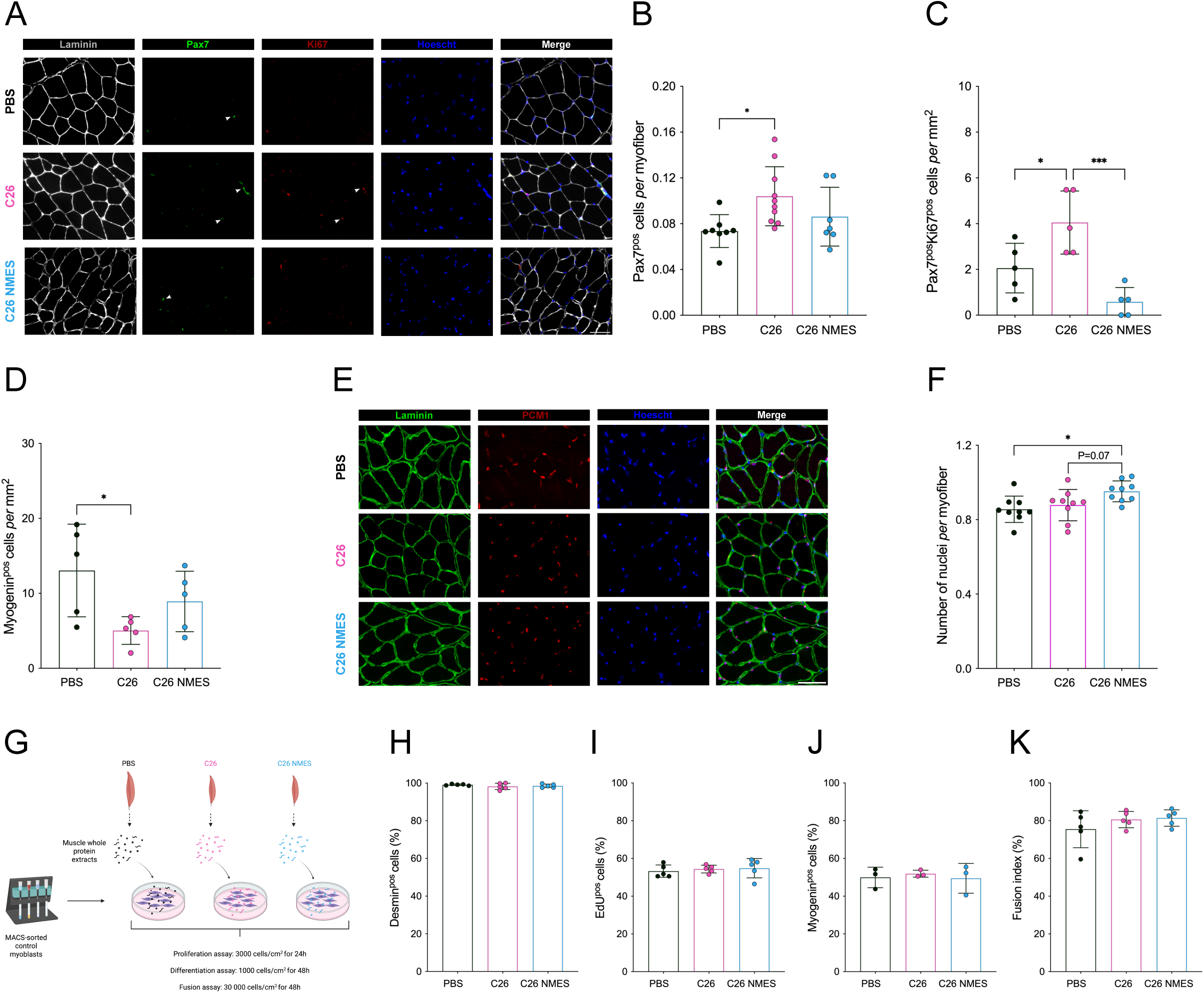
NMES training improves MuSC regulation *in vivo*. A) Immunostaining for laminin (grey), Pax7 (green), Ki67 (red) and Hoechst (blue) on *gastrocnemius* muscle section from PBS, C26 and C26 NMES mice. Scale bar: 50 µm. B) Number of Pax7^pos^ cells *per* myofiber in PBS (n=8), C26 (n=10) and C26 NMES (n=7) mice. C) Number of Pax7^pos^Ki67^pos^ cells in PBS, C26 and C26 NMES mice (n=5 *per* group). D) Number of Myogenin^pos^ cells in PBS, C26 and C26 NMES mice (n=5 *per* group). E) Immunostaining for laminin (green), PCM1 (red) and Hoechst (blue) on *gastrocnemius* muscle section from PBS, C26 and C26 NMES mice. Scale bar: 50 µm. F) Number of nuclei positive for PCM1 *per* myofiber on *gastrocnemius* muscle section from PBS (n=9), C26 (n=9) and C26 NMES (n=9) mice. G) Study design for myogenesis assays. H) Myoblast purity assessed by the percentage of desmin-positive cells. I-K) Percentage of myoblast proliferation (I), differentiation (J), and fusion index (K) in control myoblasts cultured in standard growth or differentiation medium supplemented with protein extracts (1 µg/mL) derived from PBS-, C26-, or C26 NMES-treated muscles. Each symbol represents a biological replicate. ^*,**, ***^P< 0.05, P< 0.01 and P< 0.001. Values are reported as mean ± SD.

Next, we assessed myogenin expression as a marker of MuSC differentiation. C26 tumor-bearing mice exhibited a significant reduction (−61%) in the number of myogenin-positive cells as compared to PBS-treated controls (Fig. 2D). Notably, no significant difference was observed between the PBS and C26 NMES groups.

Finally, we evaluated whether NMES influences MuSC fusion capacity by quantifying the number of PCM1-positive cells located beneath the basal lamina^35^. Consistent with our previous observation in healthy mice^33^, NMES promoted myonuclear accretion, as evidenced by a higher number of PCM1^pos^ cells *per* myofiber in C26 NMES mice (+8-11%) as compared to both C26 (P=0.07) and PBS control groups (Fig. 2E-F). Overall, these findings suggest that NMES improves MuSC fusion in C26 tumor-bearing mice.

### NMES does not affect MuSC regulation *in vitro*

Next, we aimed at investigating whether NMES affects the intrinsic properties of MuSCs *in vitro*. To address this question, we initially attempted to isolate MuSCs from PBS-treated, C26 and C26 NMES-stimulated muscles. However, since NMES was applied only to the right plantar flexor muscles, the number of MuSCs sorted from the *gastrocnemius* muscle was insufficient for myogenesis assays. To overcome these limitations, we adopted an alternative experimental approach. Myoblasts, previously isolated from healthy BALB/c mice, were cultured in media supplemented with 1 µg/mL of muscle protein extracts derived from PBS, C26 or C26 NMES-stimulated mice (Fig. 2G). Assays were performed with a cell culture purity above 95% in all conditions (Fig. 2H). MuSC proliferation was assessed by quantifying the proportion of EdU-positive cells, which showed no significant difference among the three groups (Fig. 2I). Myoblast differentiation and fusion were assessed after 48 hours of treatment with protein extracts. Muscle samples from both C26 and C26 NMES did not alter myoblast differentiation or fusion relative to PBS-treated muscle extracts (Fig 2J-K). Overall, these findings indicate that, in the absence of circulating tumor-derived factors, cachectic muscle extracts do not impair myogenesis. Consequently, when myogenic function is intact, NMES does not directly affect MuSC regulation *in vitro*.

### NMES reduces neutrophil infiltration while promoting macrophage abundance and a switch toward an anti-inflammatory profile *in vitro*

The next step was to determine whether NMES influenced other cell types of the MuSC niche, which are known to interact with MuSCs and contribute to their regulation^9^. We initially quantified the number of FAPs (PDGFRα^pos^ cells) using immunostaining, given that FAP accumulation has been previously reported in CC^14^. However, we observed no significant difference in FAP numbers among the three groups (Fig. S2).

CC is characterized by muscle inflammation as illustrated by the accumulation of immune cells, including macrophages and neutrophils, in cachectic muscle tissue^18–21^. We recently showed that NMES increases macrophage content in healthy muscles^33^. In this context, flow cytometry was used to characterize immune cell abundance and status (Fig. 3A) using a gating strategy identifying total neutrophils (CD64^neg^Ly6C^pos^) and macrophages (CD64^pos^) among leukocytes (Fig. S3). Neutrophils were reduced by 42% in the C26 NMES group as compared to C26 controls (Fig. 3A-B). Additionally, the proportion of macrophages, defined by CD64 expression, was 35% higher in C26 NMES as compared to C26 (Fig. 3A&C). We next assessed the phenotypic profile of macrophages using various cell surface markers. Both C26 and C26 NMES mice exhibited similarly low proportions of Ly6C^pos^ (<5%; Fig. 3A&D) and CCR2^pos^ macrophages (∼25%) (Fig. 3E). In contrast, Ly6C^int^ and Ly6C^neg^ macrophage populations reached 40-50% (Fig. 3A&D), with no significant differences between the two groups. These findings suggest minimal involvement of recent monocyte (CCR2^pos^) recruitment in macrophage accumulation after NMES.

**Figure 3:**
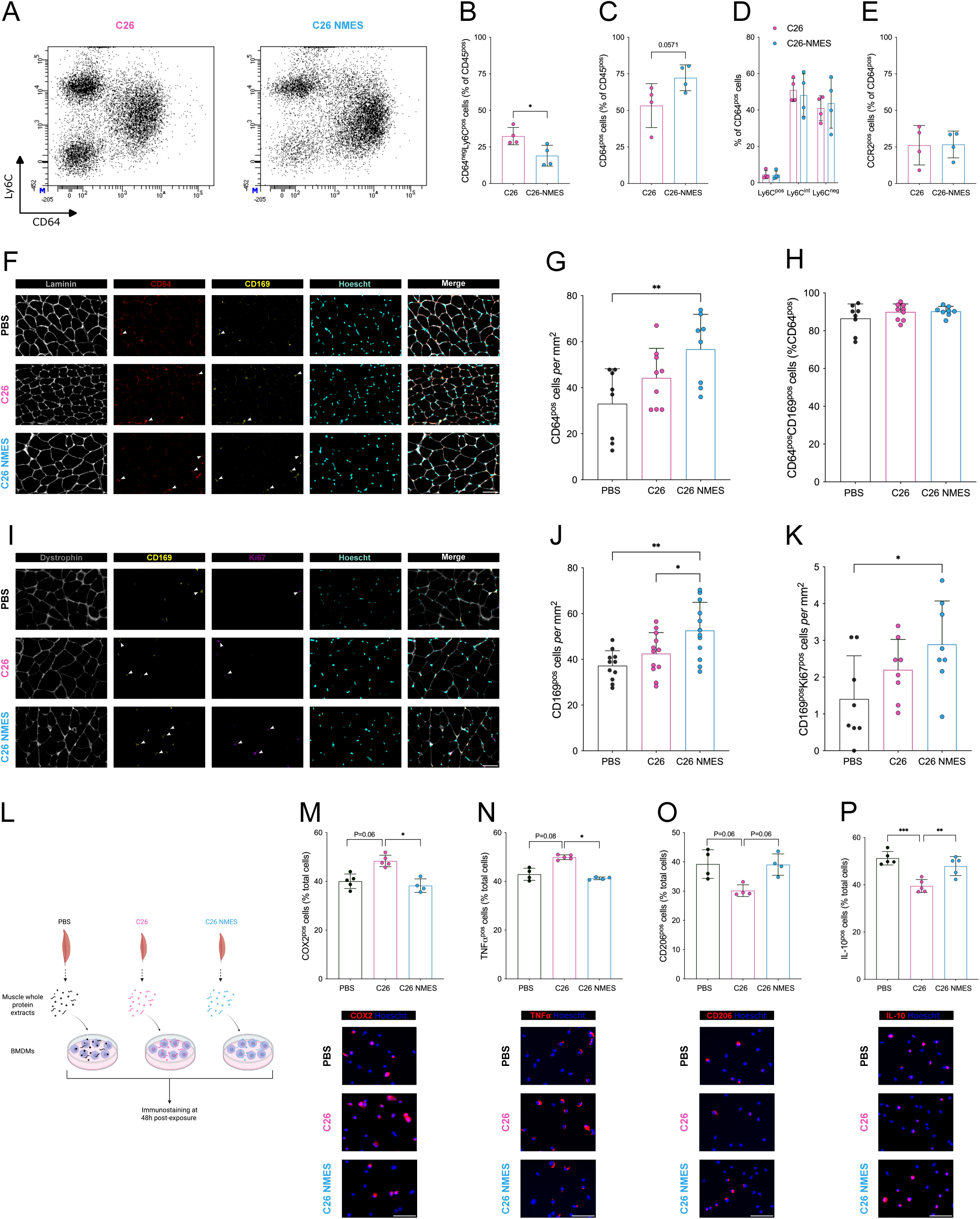
NMES reduces neutrophil infiltration, increases the number of resident macrophages and promotes a transition from a pro-towards an anti-inflammatory macrophage phenotype *in vitro* within the cachectic muscles. A) Representative dot plots. B) Proportion of neutrophils (*i.e.*, CD64^neg^Ly6C^pos^ cells) among the CD45 positive cells. C) Proportion of macrophages (*i.e.*, CD64^pos^ cells) among the CD45 positive cells. D) Proportion of macrophages (*i.e.*, CD64^pos^ cells) expressing high (Ly6C^pos^), intermediate (Ly6C^int^) or low levels of Ly6C (Ly6C^neg^). E) Proportion of macrophages expressing CCR2. Data are expressed in relation to the CD64 positive cells. For each FACS experiment, 2 to 4 muscles were included from each group (n = 4 replicates for each group). F) Immunostaining for laminin (grey), CD64 (red), CD169 (yellow) and Hoescht (cyan) on *gastrocnemius* muscle section from PBS, C26 and C26 NMES mice. Scale bar: 50 µm. G) Number of macrophages in PBS (n=8), C26 (n=9), and C26 NMES (n=8) mice. H) Proportion of macrophages positive for CD169 in PBS (n=8), C26 (n=9), and C26 NMES (n=8) mice. I) Immunostaining for dystrophin (grey), CD169 (yellow), Ki67 (purple) and Hoescht (cyan) on *gastrocnemius* muscle section from PBS, C26 and C26 NMES mice. Scale bar: 50 µm. J) Number of resident macrophages expressing CD169 in PBS (n=11), C26 (n=12), and C26 NMES (n=12) mice. K) Number of proliferating resident macrophages expressing CD169 in PBS (n=8), C26 (n=8), and C26 NMES (n=8) mice. L) Study design: bone-marrow derived macrophages (BMDMs) from healthy BALB/c mice were exposed to proteins extracts from PBS, C26 and C26 NMES muscles. Inflammatory status was analyzed by immunofluorescence after 48h post-exposure. M-P) Proportion of macrophages positive for COX2 (M), TNF*α* (N), CD206 (O) and Il-10 (P) and representative immunostaining. Scale bar: 50 µm. Data were obtained from 4-5 different protein extracts *per* group. ^*,**,***^P< 0.05, P< 0.01 and P< 0.001, respectively. Values are reported as mean ± SD.

To further investigate how NMES regulates the number and/or status of macrophages at the tissue level, macrophages were identified *via* immunostaining of cryosections using specific markers. In agreement with the flow cytometry analysis, the C26 NMES group exhibited a significantly higher number of CD64^pos^ macrophages (+41%) as compared to the PBS control (Fig. 3F-G). While not statistically significant, similar trends were observed when comparing the C26 and C26 NMES groups, as well as the PBS and C26 groups (Fig. 3F-G). We next utilized CD169, a marker that has been previously identified for labelling resident macrophages^36^. We first demonstrated that approximatively 90% of CD64^pos^ macrophages co-expressed CD169 in all conditions (Fig. 3H), confirming a minimal involvement of monocyte-derived macrophages. Interestingly, the number of CD169^pos^ macrophages was significantly higher in C26 NMES relative to the two other groups (+18-52%, Fig 3I-J). This was associated with a higher number of proliferating CD169^pos^ macrophages in C26 NMES as compared to PBS (+106%, Fig. 3I&K). Finally, we aimed at investigating whether NMES may regulate the inflammatory status of macrophages *in vitro*. Bone marrow-derived macrophages (BMDMs), from healthy BALB/c mice, were cultured in media supplemented with 1 µg/mL of muscle protein extracts derived from PBS-treated, C26 or C26 NMES-stimulated mice (Fig. 3L). BMDMs were then analyzed for the expression of various protein-level markers to assess their inflammatory status. Macrophages exposed to C26 muscle extracts showed elevated levels of the pro-inflammatory markers COX2 and TNF (+21% and +16%, respectively) as compared to PBS controls (Fig. 3M-N). Interestingly, this pro-inflammatory effect was totally abolished when macrophages were treated with extracts from C26 NMES muscles (Fig. 3M-N). Similarly, treatment with C26 muscle extracts led to a 23% reduction in the expression of the anti-inflammatory markers CD206 and IL-10, whereas NMES restored their levels to those observed in PBS-treated controls (Fig. 3O-P).

Overall, these findings suggest that NMES modulates muscle inflammation in C26 mice by limiting neutrophil infiltration, promoting the proliferation of resident macrophages *in vivo* and inducing a shift toward an anti-inflammatory phenotype *in vitro*.

## Discussion

In this study, we aimed to determine whether NMES-induced contractile activity improves muscle mass and strength and regulates MuSC fate and its niche when the training program is initiated at the stage of visible tumor development. We showed that NMES improved muscle strength and mass in the C26 mice, independently of tumor burden. These functional and structural changes were associated with improved MuSC regulation, reduced neutrophil infiltration, an increased number of resident macrophages *in vivo* and a shift from a pro-to an anti-inflammatory macrophage phenotype *in vitro* within the cachectic muscles.

Skeletal muscle functions not only as a contractile organ that generates force in response to neural stimuli, but also as an endocrine organ that releases signaling molecules. Consequently, exercise, based on repeated voluntary muscle contractions, has been proposed as a strategy to mitigate CC–induced skeletal muscle dysfunction, given its well-established benefits for muscle mass and strength in healthy individuals. To date, however, studies examining the effectiveness of exercise interventions in CC have yielded conflicting results. While some report improvements in body weight and/or muscle mass^29,30^, others observe no benefit or even detrimental effects^27,28,31^. These discrepancies may be partly explained by differences in exercise modalities, including the type of exercise (e.g., wheel running, treadmill), frequency, duration, and intensity, as well as by the timing of the intervention. Indeed, most beneficial outcomes have been reported when exercise was initiated before tumor cell inoculation or prior to tumor growth^32^, indicating that exercise was employed as a preventive rather than a therapeutic strategy against CC. Moreover, voluntary exercise is often impractical for patients with advanced cancer due to poor exercise tolerance. To overcome these limitations, muscle contractile activity was artificially increased using NMES at a stage when a visible tumor was present and muscle strength had already decreased in C26 tumor bearing mice. We demonstrated that a NMES protocol, consisting of only 16 min of submaximal isometric muscle contractions (*i.e.*, at 15% of maximal strength) over a 7-day period, increased both muscle strength, mass and myofiber size, independently of tumor mass. More importantly, we demonstrated that these beneficial adaptations occurred in the absence of muscle damage. Although electrically-evoked lengthening contractions also attenuated muscle atrophy in cachectic mice^37,38^, the clinical relevance of this approach is questionable, given that the damaging effects of this contraction modality have been documented in both animal^39^ and human^40^ studies. Overall, isometric, submaximal NMES training is a safe and effective strategy to counteract muscle weakness and wasting in cachectic mice.

CC is associated with intrinsic defects of muscle contractile function as evidence by the reduced specific torque (*i.e.*, torque normalized to muscle mass)^S1^, which has been related to several factors. One proposed mechanism is the selective degradation of myosin heavy chain (MyHC) proteins, observed in both cachectic animals^S2^ and patients^S3^, which may disrupt actin-myosin interactions and lead to contractile deficits. Additional contributing factors include impaired calcium regulation or sensitivity^S4^, as well as altered cross-bridge cycling kinetics^S4,S5^, such as a decrease in strongly bound cross-bridges and/or reduced force *per* cross-bridge. In the present study, and despite its beneficial effects on muscle strength and mass, NMES fails to compensate for the decrease in specific torque induced by CC, indicating that intrinsic muscle contractile function remains impaired. This may be partly explained by the short therapeutic window (7 days) between the development of a visible tumor and the attainment of ethical endpoints. Although evidence is limited, longer NMES training durations (>4 weeks) may improve myofiber contractility. For example, a single case study in a healthy man showed a significant increase in specific torque of type I myofibers^S6^, while 8 weeks of NMES led to increased contraction velocity of type I myofibers in patients with breast cancer undergoing chemotherapy^S7^. Future studies using genetically engineered mouse models of cachexia^S8^ could investigate the effects of NMES on contractile dysfunction over longer time windows.

Growing evidence indicates alterations in MuSC regulation in both cachectic animals^11–16^ and patients^11,17^. Indeed, tumor-derived circulating factors have been shown to disturb the MuSC myogenic program by promoting MuSC activation while impairing their differentiation and fusion due to a sustained expression of NF-kB, thereby contributing to muscle atrophy^11^. Consistent with these observations, we found that C26 tumor-bearing mice exhibited an increased number of proliferating MuSCs alongside a reduced number of myogenin-positive cells. Interestingly, NMES not only prevented this abnormal MuSC proliferation but also restored the number of myogenin-positive cells toward control levels. Moreover, NMES enhanced MuSC fusion, as evidenced by an increased number of myonuclei. These findings highlight the critical role of contractile activity in positively regulating MuSC function despite tumor-derived circulating factors. Importantly, the effects of contractile activity on MuSCs appear to be context-dependent. In our previous study on healthy mice, NMES increased the number of proliferating MuSCs^33^, whereas in the present study of cachectic mice, it reduced MuSC proliferation. This is consistent with a previous report showing that voluntary exercise prevents CC-induced Pax7 overexpression^S9^. However, exercise in that study was initiated at the time of tumor inoculation, suggesting a preventive rather than a therapeutic effect on MuSC regulation. Finally, we demonstrated that the beneficial effects of NMES on MuSC regulation occur only in the presence of cachectic factors, as our *in vitro* experiments failed to reproduce the *in vivo* findings. Indeed, myogenesis remained unaltered despite exposure to a cachectic muscle environment, consistent with previous reports describing the deleterious effects of tumor-associated factors on this process^11,13,14^. In this context, the absence of effects from muscle protein extracts derived from C26 NMES mice on myogenesis is not unexpected. These results suggest that the positive impact of NMES on MuSC regulation in cachectic mice may depend on the presence of circulating tumor-derived factors and/or on the preservation of the MuSC niche, which could be required for sensing the contractile activity generated by myofiber.

Skeletal muscle tissue inflammation has recently been described as a hallmark feature of CC. For instance, accumulation of neutrophils has been observed in several animal models of CC^20,21,S10^. Here, we reported that NMES significantly reduces the proportion of neutrophils, which may contribute to the better preservation of muscle mass in NMES-trained mice. Indeed, pharmacological depletion of neutrophils using an anti-Ly6G antibody partially or completely prevents muscle atrophy in cachectic mice, independent of tumor burden^20,S10^. However, a direct causal relationship between neutrophil modulation and the regulation of muscle mass has yet to be established. Among the immune cells accumulating in cachectic muscle, macrophages also play a pivotal role in driving muscle loss. Consistent with previous animal and clinical studies^18,19,21^, C26 tumor-bearing mice exhibited macrophage accumulation, with only a small proportion of cells expressing high levels of Ly6C and CCR2, suggesting minimal contribution from recent monocyte recruitment. In parallel, the number of CD169-positive cells, previously identified as markers of resident macrophages^36^, was increased as compared with controls. This aligns with recent findings showing that macrophages accumulating in cachectic muscle largely originate from a tissue-resident pool identified by the low expression of Ly6C^21^. Interestingly, and in agreement with our previous observations in healthy mice^33^, NMES increased the total macrophage number, through proliferation of resident macrophages, identified as Ki67- and CD169-positive cells. Although depletion of resident macrophages in cachectic muscle reduced myofiber size^21^, it also enhanced MuSC fusion, so the extent to which NMES-induced expansion of resident macrophages contributes to its beneficial effects on muscle force, mass, and MuSC regulation remains unclear. It is also noteworthy that CC is driven, at least in part, by complex interactions among macrophages, myofibers, MuSCs, and FAPs *via* NF-κB signaling^21^. Our *in vitro* experiments using BMDMs treated with muscle protein extracts from NMES-trained mice demonstrated a potent effect on inflammatory status, shifting macrophages toward an anti-inflammatory profile *in vitro*. This shift may help to normalize the disrupted cellular interactions within the MuSC niche. Overall, further studies are required to determine how NMES-induced muscle contractions modulate these complex cellular interactions and muscle inflammation in cachectic mice, and to identify the underlying molecular effectors.

Our findings may have important clinical implications. Although NMES is safe, does not require active patient cooperation, and can be self-administered at home or at the bedside, its effectiveness in cancer patients remains equivocal^S11^. This is largely due to methodological limitations, as the force generated in response to stimulation, the main determinant of NMES efficacy^S12^, has never been accurately quantified in these patients. Here, we demonstrated that NMES applied at a submaximal force level (*i.e.*, 15% of maximal force) preserves muscle force and mass. This level was chosen to reflect NMES application in severely impaired patients^S13^, for whom higher stimulation intensities are often intolerable due to discomfort associated with electrical stimuli^S12^.

In conclusion, we provide compelling evidence that NMES-induced a mild contractile activity is a potent stimulus for preserving muscle strength and mass, improving MuSC regulation and modulating muscle tissue inflammation in a mouse model of CC.

## Acknowledgments

We thank Paolo Costelli (University of Torino) for providing the C26 cell line. We thank the personnel from the Aniphy mouse facility for animal care.

## Conflict of interest

The authors disclose no competing interest

## Fundings

JG was supported by Oncostarter Cancéropôle Lyon Auvergne Rhône-Alpes CLARA, the Fondation ARC pour la Recherche sur le Cancer and the Ligue contre le Cancer (Comité de l’Isère).

## Notes

### Competing Interest Statement

The authors have declared no competing interest.

