## Supplemental materials for "NEUROMUSCULAR ELECTRICAL STIMULATION LIMITS MUSCLE WEAKNESS, ATROPHY, MODULATES SATELLITE CELL FUNCTION AND REDUCES INFLAMMATION IN CANCER CACHEXIA"

### Extended method section

#### Ethics statement

All of the experiments and procedures were conducted in accordance with French and European legislations on animal experimentation and the ARRIVE guidelines and approved by the ethical committee CEEA-55 and the French ministry of research (APAFIS#12794).

#### Torque data analysis

Torque data was sampled at 1000 Hz with a PowerLab8/35 (ADinstruments, Sydney, Australia) and analyzed with LabChart software (v8.1.17 ADInstruments, Sydney, Australia).

#### Immunostaining

Cryosections were permeabilized in Triton-X100 0.5% for 10 min at room temperature, washed 3 times in PBS and then blocked in 4% BSA for 1 hour at room temperature. Cryosections were then incubated with primary antibodies overnight 4°C, washed 3 times with PBS and further incubated with secondary antibodies for 1 hour at 37°C. The following primary antibodies were used anti-Laminin (1:200, rabbit, L9393, Sigma), anti-Laminin α2 (4H8-2) (1:1000, rat, sc-59854, Santa Cruz Biotech), anti-Myogenin (1:500, rabbit, Ab124800, Abcam), anti-PCM1 (1:1000, rabbit, HPA023370, Sigma), anti-CD64 (1:200, rat, 161002, Biolegend), anti-CD169 (1:100, rat, 142401, Biolegend) and anti-PDGFRα (1:200, goat, AF1062, R&D Systems). Secondary antibodies were: FITC AffinePure Donkey anti-rat (1:200, ref: 712-095-153), FITC AffinePure Donkey anti-rabbit (1:200, ref: 711–095–152), Cy3 AffinePure Donkey anti-rat (1:200, ref: 712–165-153), Cy3 AffinePure Donkey anti-goat (1:200, 705-165-147), Cy5 AffinePure Donkey anti-rabbit (1:200, ref : 711-175-152), Cy5 AffinePure Donkey anti-rat (1:200, ref : 712-175-150), Cy3 AffinePure Donkey anti-rabbit (1:200, ref : 711-165-152), FITC AffinePure Donkey anti-mouse (1:200, ref : 715-095-150) and Cy3 AffinePure Donkey anti-mouse (1:200, ref: 715-165-150, for determining IgG positive myofibers) supplied from Jackson ImmunoResearch. Slides were washed with PBS, counterstained with Hoechst and mounted in fluoromount-G medium.

For Pax7/Ki67 immunostaining, cryosections were first fixed with PFA 4% for 20 min, washed 3 times in PBS, permeabilized in 100% methanol (previously cool down at −20 °C) for 6 min, washed 3 times in PBS, then immerged into citrate buffer 10 mM in 90 °C hot water bath twice 5 min, washed 3 times in PBS and blocked in BSA 4% for 2–3 h. Every step was performed at room temperature, unless indicated otherwise. Cryosections were then incubated as described above with anti-laminin (1:200, rabbit, L9393, Sigma), anti-Ki67 (1:200, sheep, AF7649, R&D) and anti-Pax7 (1:30, mouse, DSHB) in blocking buffer containing BSA 4%. Biotin-conjugated donkey anti-mouse IgG1 (1:200, #BA-2000, Vector) was used against Pax7 primary antibody for 1 h at 37 °C then DTAF-conjugated streptavidin (1:1000, 016-010-084, Jackson ImmunoResearch) was used for 1 h at 37 °C. Secondary antibodies were: Cy3 AffinePure Donkey anti-sheep (1:200, ref: 713-165-147) and Cy5 AffinePure Donkey anti-rabbit (1:200, ref: 711-175-152) supplied from Jackson ImmunoResearch.

**Murine myoblast isolation**

Myoblasts were isolated from healthy BALB/c mice. All steps were performed on ice unless indicated. The entire hindlimb muscles were dissected, minced and incubated with muscle dissociation buffer (Ham's F10 medium, 10% horse serum, 1% of penicillin-streptomycin, Collagenase II 800 U/ml, Gibco) during 1 hour at 37°C in a gently shaking hot-water bath. Muscle pieces were washed and digested further by incubating with additional 3000 U/ml of collagenase II (17101015, Gibco) and 33 U/ml of dispase (D4693, Sigma), during 30 min at 37°C with agitation. After filtration to remove muscle bones, muscle chunks and tendons, red blood cells in the cell suspension were lysed by adding ACK buffer (A1049201, Gibco). Then cells were incubated with the satellite cell isolation kit mouse (130-104-268, Miltenyi biotec) in PBS, 0.5% BSA, 2mM EDTA during 30 min at 4°C. MuSCs were collected (unlabeled cells) using magnetic-activated cell sorting consisting of passing the cells through magnetic LC columns carried by a magnetic stand. After sorting, MuSCs were cultured in growth medium (DMEM F-12 (31331093, Gibco), 20% fetal bovine serum, 2% ultroser G (15950-017, Sartorius), 1% penicillin-streptomycin) within flasks coated with 0.02% of gelatin, during 3 to 4 days for myoblast expansion.

**Muscle protein extracts**

Protein extracts were obtained from the right *gastrocnemius* muscles. Briefly, 10 sections of 30 µm were produced using the NX 50 cryostat, then sections were incubated in RIPA buffer and sonicated during 6 on-off cycles of 30-60 s. Further, tissue homogenate was incubated on a rotating wheel for 2h30 at 4°C and then centrifuged at 13,000g for 20 min at 4°C. Protein extracts concentrations were determined using the Micro BCA™ Protein Assay kit (23227, Thermo Scientific). Samples were kept at -20°C until use.

**Myogenesis assessment of MuSC directly exposed to muscle protein extracts**

Myoblasts from control BALB/c mice were isolated by MACS and frozen in 90% FBS – 10% DMSO. For proliferation assay, myoblasts were thawed, counted and seeded directly at 3000 cells/cm$^2$ in a 48 well plate previously coated with Matrigel Low Growth Factor (356231, Corning). Six hours later, growth medium was replaced by new growth medium containing 1 µg/ml of muscle protein extracts (from each mouse group). Twenty-four hours later, EdU (Click-iT EdU Alexa Fluor 488 Imaging kit, C10337, Thermofisher) was added to each well to the growth medium at a final concentration of 1 µg/ml and cells were further incubated for 1 hour. Cells were then washed with PBS, fixed with 4% PFA for 15 min, washed again twice with PBS and permeabilized with 0.5% Triton-X100 during 15 min. Cells were washed again before coupling of EdU with Alexa fluor-488 substrate of the Click-iT kit reaction mixture during 30 min in dark, following the manufacturer's instructions. Cells were then washed twice in PBS and blocked in 4% BSA for 1 hour. Then, myoblasts were incubated with primary antibody anti-desmin (1:200, rabbit, 32362, Abcam) overnight at 4°C. The day after, myoblasts were washed 3 times in PBS, incubated with a Cy3-conjugated Donkey Anti-Rabbit secondary antibody (1:200, 711-165-152, Jackson Immunoresearch) for 1 hour at 37°C, washed 3 times in PBS and counterstained with Hoechst.

For fusion assays, myoblasts were thawed and recovered in growth medium during 2-3 days. Myoblasts were then counted and seeded at 30 000 cells/cm$^2$ for fusion assay in a matrigel-coated 24 well plate. Six hours later, growth medium was replaced by differentiation medium containing 1 µg/ml of muscle protein extracts (from each mouse group) for 48h. Cells were washed once with PBS, fixed with 4 % PFA for 10 min, washed 3 times with PBS, permeabilized in 0.05% Triton-X100 for 10 min, washed 3 times with PBS, and then blocked in 4% BSA for 1 hour. Cells were incubated with anti-desmin primary antibody (1:200, rabbit, 32362 Abcam) overnight at 4°C. The day after, myoblasts were further washed 3 times in PBS, incubated with a Cy3-conjugated Donkey Anti-Rabbit secondary antibody (1:200, 711-165-152, Jackson Immunoresearch) during 1 hour at 37°C and washed again 3 times with PBS. Then myoblasts were counterstained with Hoechst and were ready for microscope analysis. All steps were performed at room temperature, unless indicated.

**Flow cytometry analysis**

*Gastrocnemius* muscles of the right hindlimb from C26 and C26 NMES mice (n= 2-4) were dissected to characterize macrophage subtypes. Muscles were minced in very small pieces and digested with Collagenase B 0.2% (11088815001, Roche) during 1 hour at 37°C with agitation. Collagenase activity was blocked by the addition of 5 ml DMEM:FBS (1:1). The samples were filtered on 100 and 30 µm cell strainers. Dead cells were excluded by fixable viability Ghost dye (18452S, Cell Signaling) by labeling during 15 min. After several washing steps, cell suspension was incubated with magnetic beads conjugated to anti-CD45 antibody (130-052-301, Miltenyi Biotec) for 30 min at 4°C and $CD45^{pos}$ cells (hematopoietic cells) were isolated on a MACS Separator (130-042-109, Miltenyi Biotec) using a MS Columns (130-042-201, Miltenyi Biotec). Then $CD45^{pos}$ cells were incubated with FcR Blocking Reagent (130-059-901, Miltenyi Biotec) for another 30 min at 4°C.

**Muscle injury model**

A mouse model of muscle injury was used to establish the gating strategy for flow cytometry. Mice were anesthetized in an induction chamber with 4% isoflurane, and hindlimbs were shaved prior to intramuscular injection of 50 µL cardiotoxin (Latoxan, 12 µM) into each *gastrocnemius* muscle.

**BMDM cultures**

Bone marrow-derived macrophages (BMDMs) were prepared from adult male control BALB/c mice as previously described. BMDMs were treated with protein extracts from PBS, C26 and C26 NMES muscles for 48h. BMDMs were fixed and permeabilized prior to incubation with primary antibodies (1:50 dilution): anti-COX2 (goat, Ab23672, Abcam), anti-TNF$\alpha$ (rabbit, Ab34839, Abcam), anti-CD206 (rabbit, Ab64693, Abcam) and anti-IL-10 (rabbit, Ab9969, Abcam). Detection was performed using Cy3-conjugated secondary antibodies. Nuclei were stained with Hoechst and cells were mounted in Fluoromount.

**Image capture and analysis**

***In vivo***

Ten to fifteen images were recorded from each section with an Imager Z1 Zeiss microscope at 20 x magnification connected to a CoolSNAP MYO camera or with an Axio Observer 7 (Zeiss) connected to an ORCA-Flash4.0 LT3 Digital CMOS camera. ImageJ software was used for quantitative analysis. The number of cells stained for a given marker were either divided by the number of myofibers or by the area expressed in $mm^2$ from one picture. All data analyses were conducted in a blinded manner with respect to experimental conditions

For whole cryosection analysis, slides were automatically scanned at 10 x magnification using the Axio Observer.Z1 (Zeiss) connected to CoolSNAP HQ2 CCD Camera (photometrics) or Axio Observer 7 (Zeiss) connected to an ORCA-Flash4.0 LT3 Digital CMOS camera. The image of the whole cryosection was automatically reconstituted in MetaMorph Software. The number of IgG positive myofibers (*i.e.,* based on staining with Cy3 anti-mouse) was quantified on the whole section and normalized to the total number of myofibers. Myofiber CSA was determined on the whole *gastrocnemius* muscle sections labeled by anti-laminin antibody as previously described[33,S14]. These analyses were conducted on 6715 $\pm$ 936 myofibers.

***In vitro***

For myoblasts assessment of myogenesis *in vitro*, 10 to 20 images were recorded from each well of the cell culture plate at 10 x magnification using the Axio Observer.Z1 (Zeiss) connected to CoolSNAP HQ2 CCD Camera. Quantification of $EdU^{pos}$ nuclei, $Desmin^{pos}$ cells and $Myogenin^{pos}$ nuclei was performed with ImageJ software. The percentage of proliferative myoblasts was calculated using the number of $Desmin^{pos}$ cells with $EdU^{pos}$ nuclei over the total $Desmin^{pos}$ cells. The percentage of differentiation was calculated using the number of $Myogenin^{pos}$ nuclei over the total number of nuclei. The fusion index was calculated using the number of nuclei of myotubes presenting two or more nuclei over the total number of nuclei (in both myoblasts and myotubes). Purity was assessed by counting the number of $Desmin^{pos}$ cells over the total number of cells (without desmin staining). For BMDM experiments, approximately 10 images were acquired randomly, and positive cells were quantified using ImageJ software.

#### Supplemental figure legends

**Figure S1:** A) Individual training intensity (expressed in percentage of maximal isometric torque) throughout the 80 stimulation trains of the NMES training protocol (mean of the 6 sessions). Each symbol represents an individual mouse (n=40 mice). For the sake of clarity, values are reported as mean $\pm$ SEM. B) Proportion of myofibers positive for IgG in PBS (n=3), C26 (n=7) and C26 NMES (n=8). Values are reported as mean $\pm$ SD.

**Figure S2:** Number of fibro-adipogenic progenitors (PDGFR$\alpha^{pos}$ cells) in PBS (n=7), C26 (n=8) and C26 NMES (n=9).

**Figure S3:** Flow cytometry analysis of the various cell types isolated from the *gastrocnemius* muscle in C26 and C26 NMES mice. Cells were separated by magnetic beads based on their CD45 expression level. CD45$^{pos}$ cells were incubated with Fc block and labelled with CD45, CD64, and Ly6C antibodies to isolate neutrophils (Neut) and macrophages (*i.e.*, CD64$^{pos}$ cells; Macro.) expressing high (Ly6C$^{pos}$), intermediate (Ly6C$^{int}$) or low levels of Ly6C (Ly6C$^{neg}$). The gating strategy used to identify CCR2-positive macrophages is also shown (bottom right panel). For each experiment, 2 to 4 muscles were digested from each group.

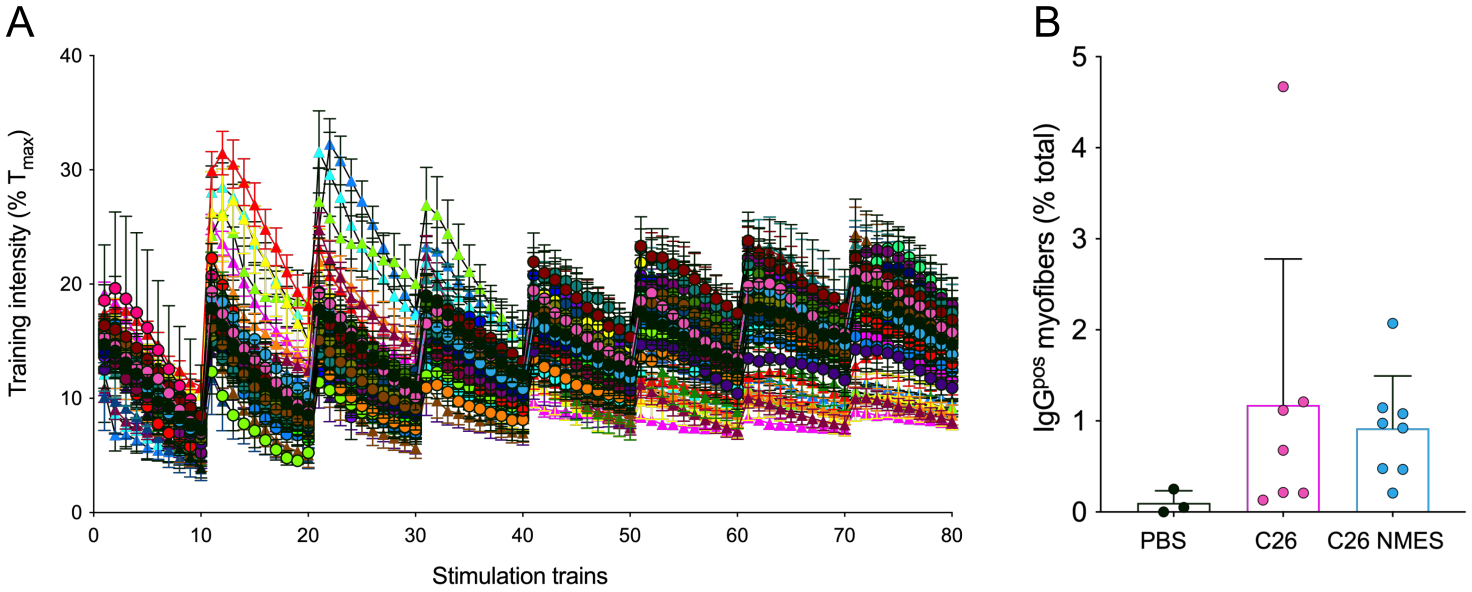


Supplemental Figure 1

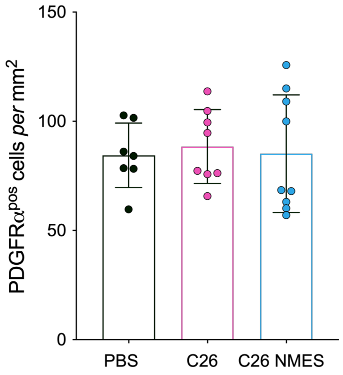


Supplemental Figure 2

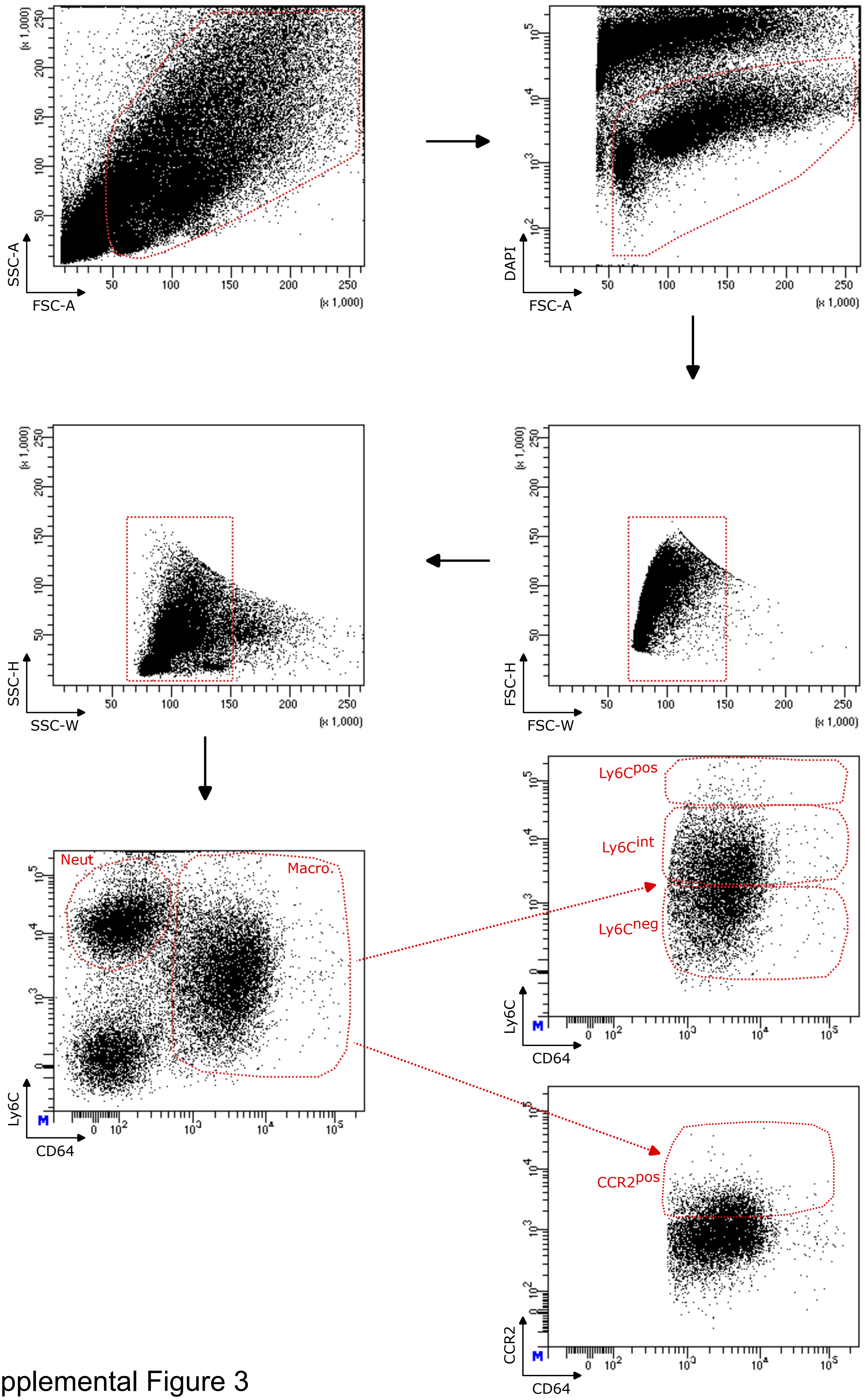


Supplemental Figure 3
